# Meiotic viral attenuation through an ancestral apoptotic pathway

**DOI:** 10.1101/521237

**Authors:** Jie Gao, Sabrina Chau, Fuad Chowdhury, Tina Zhou, Saif Hossain, G. Angus McQuibban, Marc D. Meneghini

## Abstract

The programmed release of apoptogenic proteins from mitochondria is a core event of apoptosis, though ancestral roles of this phenomenon are not known. In mammals, one such apoptogenic protein is Endonuclease G (EndoG), a conserved nuclease that fragments the DNA of dying cells. In this work, we show that budding yeast executes meiotically programmed mitochondrial release of an EndoG homologue, Nuc1, during sporulation. In contrast to EndoG’s ostensible pro-death function during apoptosis, Nuc1 mitochondrial release attenuates the cytosolic dsRNA mycovirus, Killer, protecting spores from a lethal accumulation of its encoded toxin. Our identification of cell-protective viral attenuation as a target of this rudimentary apoptotic pathway illuminates a primordial role for mitochondrial release of EndoG.

**One Sentence Summary:** Yeast sporulation induces release of mitochondrial endonuclease G to accomplish viral attenuation.

## Main Text

Mathematical models suggest that host-virus conflicts drove the evolution of programmed cell death (PCD) in single celled eukaryotes*(1, 2)*. Unicellular yeast species harbor numerous cytosolic viruses that have no extracellular phase and are only vertically transmitted through cytoplasmic inheritance*(3)*. The most comprehensively studied of these is Killer, a paired system of the L-A and M double stranded RNA (dsRNA) viruses in *Saccharomyces cerevisiae.* L-A produces a viral particle that houses and propagates M, which itself encodes a secreted toxin that kills neighboring uninfected cells and confers immunity to the host*(3)*. Genetic studies reveal that Killer exists in clear conflict with its host, though how and if this relates to PCD or other developmental occurrences in yeast is not known*(4)*.

Yeast sporulation employs internal meiotic divisions and culminates in the development of spore progeny within the remnant of the mother cell*(5)*. PCD of this remnant cell occurs as an intrinsic aspect of sporulation and is executed through vacuolar rupture, leading to mother cell autolysis*(6, 7)*. Spores survive this process, called meiotic PCD, through coordinated development of their protective spore coats*(6)*. Under conditions of reduced carbon, undeveloped meiotic nuclei are frequently swept up in meiotic PCD and their DNA is fragmented into nucleosomal ladders in a manner requiring *NUC1,* the yeast homolog of the EndoG family of mitochondrial nucleases*(7)*. This finding is reminiscent of EndoG promoting DNA fragmentation of apoptotic cells following its mitochondrial release, though

EndoG is not required for apoptosis *per se* and its roles in this process are debated*(8–12)*. *NUC1* function is similarly puzzling, as meiotic PCD is unperturbed in *nuc1Δ/Δ* mutants beyond the absence of fragmented DNA*(7)*. Indeed, although activated nuclease pathways that promote genome fragmentation accompany diverse forms of PCD in plants, animals, protists, and fungi, the adaptive purposes for this association are unclear*(13–15)*.

The observation of NUC1-dependent DNA ladders that can accompany meiotic PCD suggests that Nuc1 is released from mitochondria during sporulation. We confirmed Nuc1 mitochondrial localization during meiosis using fluorescence microscopy of Nuc1-GFP and mito-RFP expressing cells in the efficiently sporulating SK1 strain background, which accomplishes meiotic nuclear divisions in 8-10 hours*(7)*(**Fig. 1A**). Interestingly beginning at the 6-hour timepoint, some Nuc1-GFP signal was detected that did not appear to overlap with mitochondria (**Fig. 1A**). To more sensitively assess the sub-cellular accumulation of Nuc1, we measured the abundance of a FLAG epitope tagged allele of Nuc1 in biochemically prepared cytosolic and mitochondrial fractions from cells undergoing sporulation. The integrity of these cell compartment fractions was first confirmed using western blotting to detect proteins known to localize to the mitochondria or the cytosol (**Fig. S1**). As expected, Nuc1-FLAG fractionated exclusively with mitochondria at the onset of sporulation (**Fig. 1B**).

**Figure 1.**
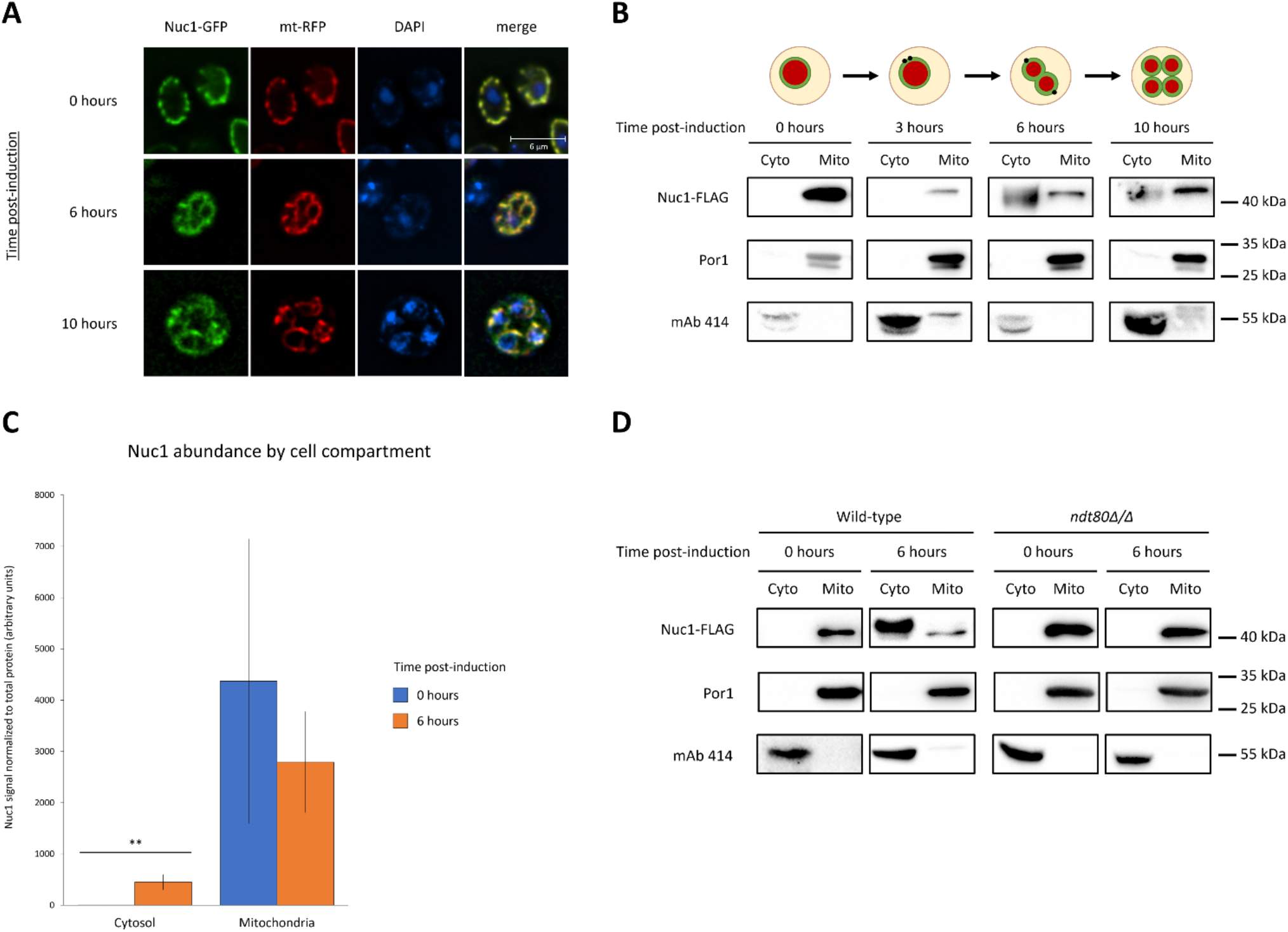
Developmental regulation of Nuc1 location in meiosis. **(A)** Nuc1 localization assessed at 0, 6, and 10 hours post-induction of sporulation in fixed cells expressing Nuc1-GFP and mito-RFP (mt-RFP). Cells were stained with DAPI. Scale bar, 6 μm. **B**) Anti-FLAG, anti-Por1, and mAb 414 Western blots of mitochondrial (mito) and concentrated cytosolic (cyto) cell fractions sampled from a synchronized meiotic timecourse of a strain expressing Nuc1-FLAG. A schematic depiction of meiotic progression at each timepoint is shown above the corresponding Western blots. Fractions were probed with antibodies against Por1 and mAb 414 to validate successful enrichment of mitochondrial and cytosolic contents, respectively. Molecular weight markers corresponding to blots are given on the far right. **(C)** Quantification of Nuc1 abundance in cytosolic and mitochondrial cell fractions at 0 and 6 hours post-induction (n = 4 per timepoint). Data points represent mean ± standard deviation. **Difference in Nuc1 abundance is statistically significant (two-tailed T-test, *p* < 0.01). **(D)** Anti-FLAG Western blots of cell fractions sampled at meiotic timepoints from wild-type and a mutant *ndt80Δ/Δ* strain expressing Nuc1-FLAG (fractionation controls included). Molecular weight markers on the far right.

Beginning at the 6-hour timepoint and persisting through the post-meiotic 10-hour timepoint, we detected robust Nuc1-FLAG accumulation in concentrated cytosolic fractions (**Fig. 1B**). We quantified this in 4 biological replicates and determined that a small but significant fraction of total Nuc1 (13%) accumulated in the cytosol during meiosis, suggesting that a portion of Nuc1 was released from mitochondria (**Fig. 1C**). *NDT80* encodes a transcription factor that commits cells to sporulation and directs meiotic PCD*(5, 6, 16). NDT80* was required for Nuc1-FLAG mitochondrial release, confirming this occurrence as an intrinsic and NDT80-programmed aspect of sporulation (**Fig. 1D**).

Our fractionation experiments were performed using standard sporulation conditions that do not exhibit NUC1-dependent DNA ladders from uncellularized meiotic nuclei*(7)*. We therefore considered other roles for cytosolic Nuc1, which, like EndoG, can digest RNA as well as DNA*(17, 18)*. Consistent with an early study*(19)*, we found that a *nuc1Δ/Δ* diploid strain exhibited increased abundance of L-A Gag, the major coat protein of the L-A viral particle (**Fig. 2A**). Curiously, L-A Gag levels plummeted following sporulation in both wild-type and *nuc1Δ/Δ* strains (**Fig. 2A**). We were unable to detect Killer toxin within wild-type cells prior to or following sporulation, reflecting its known secretion into the cell supernatant (**Fig. 2A**)*(20, 21)*. In striking contrast, we found that the sporulated *nuc1Δ/Δ* strain accumulated substantial amounts of Killer toxin (**Fig. 2A**). These findings reveal that *NUC1* attenuated M expression specifically during sporulation. We further investigated this using RT-qPCR and found that while bulk M RNA levels were repressed 5-fold during wild-type sporulation, they were only repressed 3.7-fold following sporulation of a *nuc1 Δ/Δ* strain (**Fig. S2**). As it is not possible to distinguish the Killer toxin-encoding transcript from its dsRNA genome, this subtle effect of *nuc1 Δ/Δ* may reflect a major role for Nuc1 in M transcript repression that is largely masked from detection by the dsRNA M genome. This interpretation is consistent with our western blot experiments as well as Nuc1’s ability to digest ssRNA but not dsRNA*(17)*.

**Figure 2.**
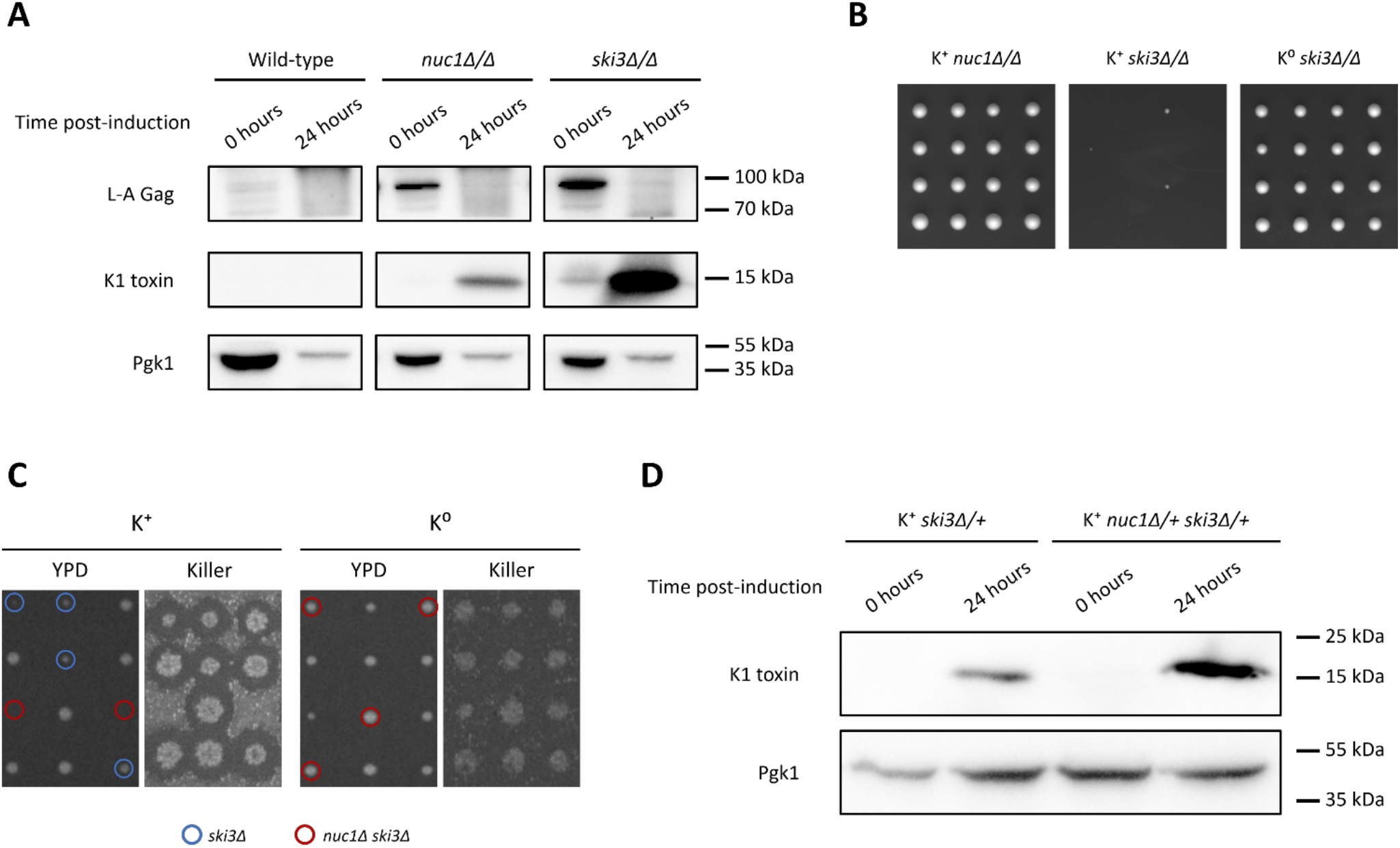
Nuc1 mediates viral repression in meiotic yeast. **(A)** Anti-Killer toxin and anti-L-A gag Western blots of whole-cell lysates sampled from synchronized sporulations of homozygous K+ strains at 0 and 24 hours post-induction of sporulation with Pgk1 loading control. Molecular weight markers are shown on the right. **(B)** Subsection of homozygous mutant dissections with 4 tetrads per strain, 3 days post-dissection on YPD at 30°C. **(C)** Tetrad dissections of K+ (left) and K^0^ (right) *nuc1Δ/+ ski3Δ/+* mutants. Left panels, 2 days post-dissection on YPD at 30°C; right panels, Killer plates showing killing activity. Known or inferred haploid genotypes of interest are circled in blue *(ski3Δ,* K+ dissection only) and red *(nuc1Δ ski3Δ).* **(D)** Anti-Killer toxin Western blots of whole-cell lysates sampled from synchronized sporulations of heterozygous K+ strains at 0 and 24 hours post-induction of sporulation with Pgk1 loading control. Molecular weight markers are shown on the right.

Our findings illuminate a sporulation-specific role of *NUC1* in Killer attenuation and we therefore considered the roles of established anti-Killer pathways during sporulation. The *SKI2, 3,* and *8 (<u>s</u>uper<u>ki</u>ller)* genes encode subunits of a well-characterized cytoplasmic complex that attenuates Killer expression in vegetative cells through translational repression of M-encoded transcripts*(3, 22)*. Consistent with a previous study, we found that deletion of both copies of *SKI3* caused increased L-A Gag levels in mitotic cells (**Fig. 2A**)*(4)*. As with what we observed in *NUC1* mutants, detectable Gag was subsequently squelched during sporulation of a *ski3Δ/Δ* strain (**Fig. 2A**). Sporulation of homozygous *SKI3* mutants caused a large accumulation of Killer toxin and an according defect in bulk M RNA repression (**Fig. 2A and S2**). While *nuc1 Δ/Δ* and *ski3Δ/Δ* both sporulated efficiently, *ski3Δ/Δ* produced nearly 100% inviable spores, suggesting that the high degree of Killer toxin accumulation within them caused their lethality (**Fig. 2B**). To confirm this, we generated a *ski3*Δ/Δ K^0^ strain that lacked the M virus and found that Killer loss completely suppressed the lethal phenotype (**Fig. 2B**). These findings identify a heretofore-unknown essential role for Killer attenuation during sporulation that is dependent on the SKI complex, and show that *NUC1* contributes to this attenuation.

To assess the relative contributions of *NUC1* and *SKI3,* we dissected a doubly-heterozygous *nuc1Δ/+ ski3Δ/+* diploid strain and found that *nuc1 Δ ski3Δ* spores were inviable (**Fig. 2C**). Sporulation of this *nuc1Δ/+ ski3Δ/+* strain caused an accumulation of Killer toxin that was substantially greater than that exhibited by a *ski3Δ/+* strain, suggesting that elevated Killer toxin levels in *nuc1 Δ ski3Δ* spores accounted for their lethality (**Fig. 2D**). Indeed, eliminating the M virus fully suppressed *nuc1 Δ ski3Δ* synthetic lethality, confirming that *NUC1* and *SKI3* attenuated lethal Killer toxin accumulation during sporulation through parallel mechanisms (**Fig. 2C**). To determine if the nuclease activity of Nuc1 was required for Killer attenuation, we employed a previously characterized enzymatically dead allele of Nuc1, Nuc1-H138A*(23)*. We introduced plasmids expressing either *NUC1* or *NUC1(H138A)* under the control of the inducible *GAL1-10* promoter into a *nuc1Δ/+ ski3*/+* diploid strain. We found that over-expression of *NUC1* during spore germination, but not of *NUC1(H138A),* rescued the *nuc1Δ ski3Δ* synthetic lethality, suggesting that the nuclease activity of Nuc1 was essential for its attenuation of Killer virus (**Table S1**).Western blotting and cell fractionation experiments showed that Nuc1 and Nuc1(H138A) were produced to comparable levels and that some cytosolic accumulation accompanied their overexpression, consistent with attenuation of cytosolic Killer by Nuc1 (**Fig. S3**).

Notably, *ski3Δ* single mutant spore clones produced by sporulation of a *nuc1Δ/+ ski3Δ/+* strain exhibited a slight degree of slow growth, but were always viable (**Fig. 2C**). The viability of these *ski3Δ* spores contrasts with the Killer-dependent lethality exhibited by spores produced from a *ski3Δ/Δ* homozygous strain (**Fig. 2B**). Together, these genetic results highlight an essential role for *SKI3* in Killer attenuation during sporulation that is conferred by the diploid genome during meiosis, preceding sporogenesis and its associated spore-autonomous mode of gene expression. We refer to this as the maternal stage of sporulation.

As we observed Nuc1 mitochondrial release to initiate during maternal meiotic stages and to persist into sporogenesis (**Fig. 1B**), we devised a genetic experiment to determine the relative contributions of maternal versus spore autonomous *NUC1* function for Killer attenuation. *MAK3* encodes the enzymatic subunit of the NatC N-terminal acetyltransferase complex that is essential for Killer maintenance in vegetative cells*(24)*. Foundational studies reveal that NatC is not required for Killer maintenance during sporogenesis; rather, spore clones lacking NatC only lose Killer following germination and subsequent mitotic proliferation*(25, 26)*. We constructed a *nuc1Δ/+ ski3Δ/+ mak3Δ/+* triply heterozygous diploid strain and measured spore colony growth following its sporulation and dissection. As expected, *mak3Δ* always segregated with a Killer minus phenotype (data not shown).

The lethality of *nuc1Δ ski3Δ* mutants was rescued by *mak3Δ* in 90% of the triple mutants (**Fig. 3A**). Though the rescued spore clones showed variably delayed post-germination growth kinetics, upon subculture, the *nuc1Δ ski3Δ mak3Δ* isolates exhibited normal growth (**Fig. 3A and 3B**). These results show that the lethality caused by loss of spore-autonomous Killer attenuation by *SKI3* and *NUC1* could be overcome if the nascent colony lacked *MAK3* function. In contrast, *nuc1Δ ski3Δ mak3Δ* spores produced by dissecting a *nuc1Δ/Δ ski3Δ/+ mak3Δ/+* strain remained inviable, revealing that loss of maternal *NUC1* and spore-autonomous *SKI3* resulted in insurmountable Killer activity (**Fig. 3C**). These findings genetically identify a critical maternal role for *NUC1* in Killer attenuation, corresponding with when we first observe Nuc1 mitochondrial release.

**Figure 3.**
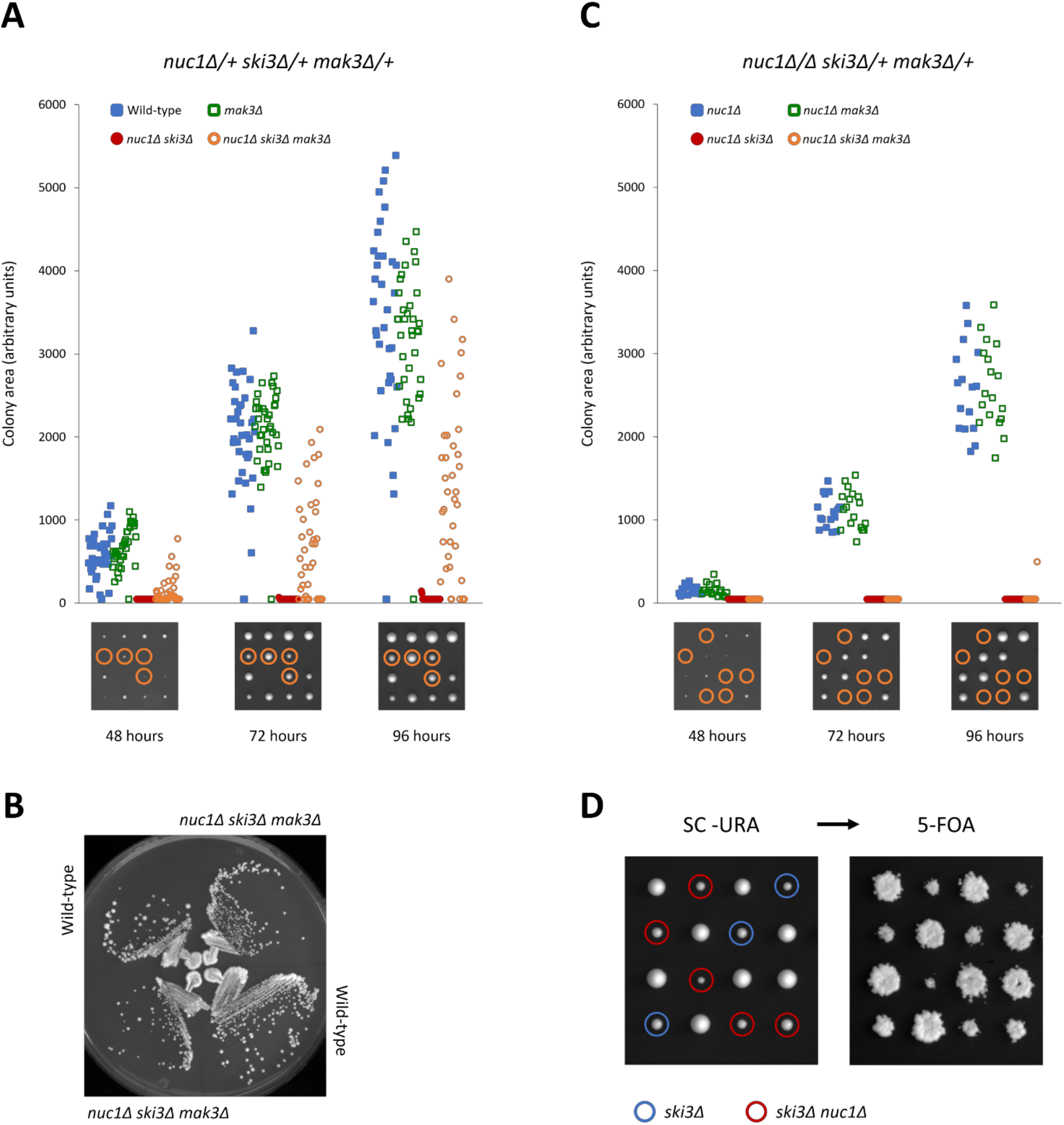
Maternal requirement of Nuc1 yeast antiviral activity. **(A and C)** Representative dissections on YPD at 30°C shown above quantifications of colony growth at the corresponding timepoints. Haploid meiotic *nuc1Δ ski3Δ mak3Δ* progeny are circled. (Bottom) Growth plots of known/inferred haploid genotypes of interest were derived by measuring colony surface area on plates imaged at each timepoint. Data sets were assembled from multiple independent dissections, with *n =* 143 colonies measured for *nuc1Δ/+ ski3Δ/+ mak3Δ/+* dissections and (C) *n* = 60 colonies measured for *nuc1Δ/Δ ski3Δ/+ mak3Δ*/+. **(B)** Growth of wild-type and *nuc1Δ ski3Δ mak3Δ* haploids on YPD at 30°C, imaged 2 days post streaking. **(D)** The K+ *nuc1Δ/+ ski3Δ/+* parent strain was transformed with a plasmid expressing Nuc1 from its endogenous promoter and sporulated for tetrad analysis. Plates were imaged 3 days post-dissection on SC–URA at 30°C (left), then replica-plated to SC–URA + 5-FOA (right). Genotypes of interest are denoted by blue *(ski3Δ)* and red *(nuc1Δ ski3Δ)* circles.

To confirm that *NUC1’s* essential role in Killer attenuation occurred during sporulation, we first rescued the *nuc1Δ ski3Δ* synthetic lethality by dissecting a *nuc1Δ/+ ski3Δ/+* strain containing a *NUC1-URA3* plasmid. Consistent with the hypothesis that *NUC1’s* essential function in Killer attenuation occurred during sporulation, the mitotically proliferating *nuc1Δ ski3Δ* mutants could readily lose the *NUC1* rescuing plasmid (**Figure 3D**).

We reasoned that Nuc1 mitochondrial release during sporulation might occur through a conserved mechanism. The mitochondrial voltage dependent anion channel (VDAC) functions as a pore for small hydrophilic molecules and has been implicated in the release of apoptogenic proteins, including EndoG, through the formation of an oligomeric “megachannel”*(27–30)*. Yeast VDAC is encoded by the paralogous *POR1/2* genes and ribosome profiling studies revealed that *POR2* translation is dramatically upregulated during meiosis (**Fig. S4**)*(31)*. To investigate the possibility that yeast VDAC is involved in meiotic Nuc1 release, we first determined the expression and localization of Por1/2 using a Por1-GFP Por2-mCherry expressing strain. As expected, Por1-GFP exhibited strong signal corresponding to mitochondrial networks and co-localized with mitochondrial nucleoids throughout meiosis (**Fig. 4A**). Although Por2-mCherry exhibited considerably less bright signal, it co-localized with Por1-GFP (**Fig. 4A**). Consistent with ribosome profiling studies, we found that Por2-mCherry exhibited induced expression during meiosis. However, the induced Por2-mCherry signal did not co-localize with Por1-GFP, mitochondrial nucleoids, or other mitochondrial proteins, and instead appeared as cytoplasmic foci (**Fig. 4A** and data not shown). Although Por2-mCherry localized with Por1-GFP during sporogenesis at the 10-hour timepoint, substantial amounts of Por2-mCherry persisted in the mother cell compartment (**Fig. 4A**). While we do not know the identity or significance of this Por2 structure, our findings confirm that the yeast VDAC paralogs are present on mitochondria throughout sporulation and that Por2 is meiotically induced.

**Figure 4.**
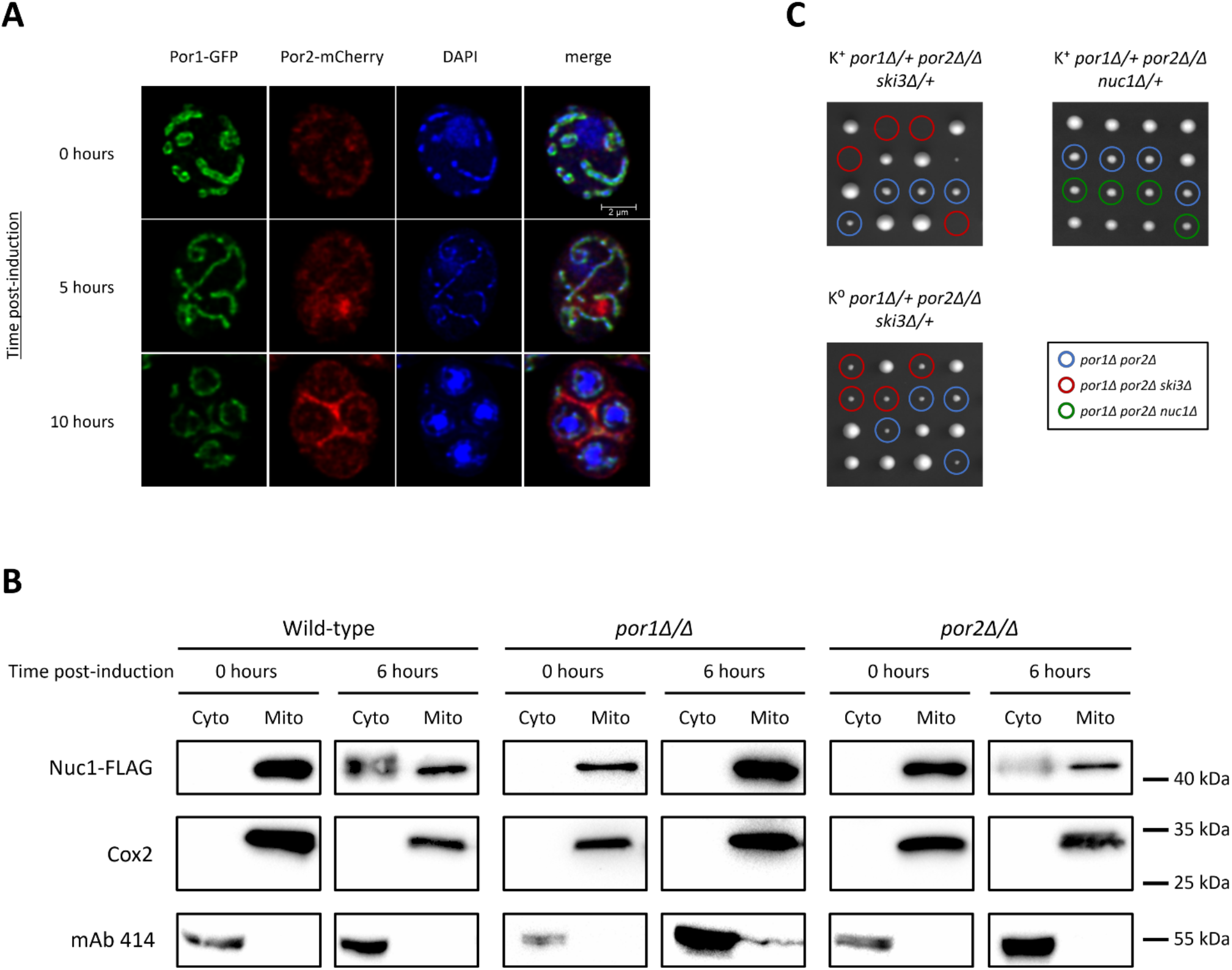
Mitochondrial VDACs mediate Nuc1 antiviral activity and cytosolic accumulation. **(A)** Meiotic imaging timecourse of cells expressing Por1-GFP and Por2-mCherry. Fixed cells were stained with DAPI to visualize nucleus and mitochondrial nucleoids. Scale bar, 2 μm. **(B)** Anti-FLAG Western blots of cell fractions sampled from a synchronized sporulation of VDAC mutants expressing Nuc1-FLAG with a wild-type strain for comparison (fractionation controls included). Note that the *por2Δ/Δ* and *por1Δ/Δ* cytosolic lanes were loaded with twice the sample volume of wild-type. Molecular weight markers are shown on the right. **(C)** Dissections of *VDAC-NUC1/SKI3* triple mutants, imaged 3 days post-dissection on YPD at 30°C. Known/inferred haploid genotypes of interest are indicated: *por1Δ por2Δ* (blue), *por1Δ por2Δ ski3Δ* (red), and *por1Δ por2Δ nuc1Δ* (green).

We next assessed the roles of yeast VDAC in Nuc1 mitochondrial release using subcellular fractionations. Nuc1-FLAG was undetectable and weakly detectable respectively in cytosolic preparations of sporulating *por1Δ/Δ* or *por2Δ/Δ* strains at the 6-hour meiotic timepoint (**Fig. 4B**). These results suggest that Nuc1 undergoes programmed mitochondrial release during meiosis through a conserved VDAC mechanism.

A parsimonious model explaining our findings posits that Por1/2 VDAC-mediated release of Nuc1 into the cytosol exposes M transcripts to Nuc1, resulting in Killer attenuation. A key prediction of this model is that yeast VDAC should act in a *NUC1* antiviral genetic pathway. As deletion of both copies of *POR1* caused a strong defect in Nuc1 mitochondrial release at the 6-hour maternal meiotic stage, we first tested this prediction by sporulating and dissecting a *por1Δ/Δ ski3Δ/+* diploid strain. Double mutant spores lacking *SKI3* produced by these strains exhibited a high degree of lethality, and were recovered at only 1/3 the expected frequency (**Table S2**). Consistent with our model, this partially penetrant synthetic lethality was fully suppressed by loss of Killer and not observed in *por1Δ/Δ nuc1Δ/+* dissections (**Table S2**). We next investigated the contributions of *POR2* to the putative Por1/2-Nuc1 antiviral pathway. Following sporulation of a *por1Δ/+ por2Δ/Δ ski3Δ/+* strain, we found near 100% lethality of *por1Δ por2Δ ski3Δ* mutant spores (**Fig. 4C and Table S2**). Though *por1Δ por2Δ* spore clones were slow growing, the lethality of *por1Δ por2Δ ski3Δ* mutant spores was fully suppressed by loss of Killer (**Fig. 4C and Table S2**). Dissection of a*por1Δ/+ por2Δ/Δ nuc1Δ/+* strain resulted in fully viable *por1Δ por2Δ nuc1Δ* spores, confirming that Por1/2 acted in a Nuc1 pathway to attenuate Killer during sporulation (**Fig. 4C and Table S2**).

Like with *NUC1, POR1* mutation has been associated with increased L-A abundance in haploid cells following saturating growth and media exhaustion*(19, 32)*. It thus seems likely that the Por1/2-Nuc1 pathway we have identified can function outside of meiosis, though our results show its meiotic role is much more consequential. It is important to note that undomesticated *S. cerevisiae* exist essentially exclusively as diploids, and that the response of these diploid cells to media exhaustion is to sporulate*(7)*. Vestigial applications of the Por1/2-Nuc1 meiotic pathway in artificially neutered haploid cells may thus be expected to occur in response to these similar nutritional cues. More broadly, much of the controversial phenomenon of yeast “apoptosis” observed to occur in haploid cells following their exhaustion of growth media may relate to vestigial manifestations of meiotic PCD*(33)*.

Although altruistic scenarios have been invoked to answer the paradox “why would a unicellular organism ever commit suicide?”, transferring the paradigm of altruistic PCD from multicellular contexts to unicellular ones remains dubious and may cloud our understanding of the nature and origin of microbial PCD *(34–36).* Our findings identify meiosis as a context where crucial host-virus interactions are mediated within the milieu of PCD in budding yeast. While the Por1/2-Nuc1 pathway is similar to VDAC-EndoG pathways that accompany canonical apoptosis, rather than death, Por1/2-Nuc1 promotes survival. As cell death pathways must have arisen from non-suicidal roles, our findings may thus illuminate transitional forms of PCD.

## Acknowledgements

We thank Reed Wickner for strains and the L-A antibody as well as Motomasa Tanaka for the Killer antibody. We are grateful to Dr. Charlie Boone and Dr. Brenda Andrews for providing strains. Work in the Meneghini lab is supported by an NSERC Discovery grant and CIHR grant MOP-89996 to M.D.M.

## Author contributions

All of the authors contributed to conceiving the experiments. J.G., S.C., F.C., T.Z., S.H., and M.D.M. conducted the experiments and analysis. J.G. and M.D.M. wrote the manuscript.

## Data and materials availability

All data is available in the main text or the supplementary materials.

## Supplementary Materials

Materials and Methods

## Supplementary Materials for Meiotic viral attenuation through an ancestral apoptotic pathway

### MATERIALS AND METHODS

#### Strains and media

Standard *S. cerevisiae* genetic and strain manipulation techniques were used for strain construction and tetrad analysis (*1*). Refer to **Table S3** for strains used in this paper. For tetrad dissections, sporulated strains were incubated in 3 mg/mL 20T Zymolyase (Seikagaku Glycobiology) for 15-20 minutes at room temperature and spread on 1% yeast extract, 2% peptone, 0.004% adenine, 2% dextrose (YPD) plates for dissecting.

#### Plasmids

The p5472 *NUC1 CEN/ARS URA3* MOBY plasmid was obtained from Brenda Andrews (2). The pRS424 *ADH-mtRFP* plasmid was obtained from Janet Shaw (3). The pME1 *NUC1::EGFP URA3* plasmid was generated from the pUG35 *NUC1::EGFP URA3* plasmid, obtained from Frank Madeo (4), by replacement of the *MET25* promoter with the native *NUC1* promoter using SacI and XbaI (New England Biolabs). The *pESC GALpr-NUC1::FLAG HIS3* and g*pESC GALpr-NUC1-H138A::FLAG HIS3* plasmids were obtained from Frank Madeo (*4*).

#### Killer activity assay

To determine the Killer phenotypes, strains were replica-plated onto lawns of Killer-sensitive yeast cells (obtained from Reed Wickner) on buffered plates (0.0034% w/v methylene blue in YPD, adjusted to pH 4.5 with citric acid monohydrate and sodium phosphate dibasic) and grown at 20°C for 3 days to observe killer activity.

#### Anisomycin eviction of viruses

Cells were treated with 32 mM Anisomycin (BioShop ANS245) in liquid YPD for 4 days at 30°C and plated for single colonies on YPD plates to assay for Killer activity. Non-Killer isolates were confirmed to have lost the M1 genome by RT-PCR.

#### SK1 synchronized sporulation protocol

Synchronized of SK1 strains were performed using methods described by Eastwood and Meneghini (5). All sporulations are performed in standard high-carbon conditions.

#### Yeast subcellular fractionation

The Mitochondrial Yeast Isolation kit and protocol (Abcam, ab178779) was used to fractionate yeast cells by differential centrifugation. Following pelleting of the mitochondrial fraction, the soluble cytosolic fraction was collected by precipitation with 2% trichloroacetic (TCA) at 4°C for 30 minutes and washed three times with acetone before resuspension in SDS-PAGE sample buffer (50 mM Tris-HCl pH 6.8, 2% SDS, 10% glycerol, 0.025% bromophenol blue, 100 mM DTT) and heating at 100°C for 10 minutes. Mitochondrial and whole cell pellets were resuspended and heated in SDS-PAGE sample buffer.

#### Yeast whole-cell protein extraction

Vegetative cells were permeabilized with 0.1 N NaOH for 5 minutes at room temperature prior to resuspension in SDS-PAGE buffer and heating at 100°C for 10 minutes. The soluble fraction was isolated by centrifugation and used for subsequent Western blotting. Sporulated cells were disrupted by bead-beating cells suspended in SDS-PAGE buffer for 10 minutes prior to heating at 100°C.

#### Western blots

Cell fractions and whole cell lysates prepared as described above were electrophoresed on 8-10% SDS-PAGE and transferred to PVDF membranes. Blots were incubated overnight in diluted primary antibody, probed with horseradish peroxidase (HRP)-conjugated horse anti-mouse (Cell Signaling Technology #7075) or goat anti-rabbit (Cell Signalling Technology #7074) secondary antibody, and detected using Luminata Forte Western HRP Substrate (EMD Millipore). Blots were imaged with the Bio-Rad ChemiDoc XRS+ system; image processing was performed with the Image Lab software package (Bio-Rad). The primary antibodies and dilutions used were 1:1000 anti-FLAG M2 (Sigma-Aldrich cat# F1804), 1:2000 mouse anti-nuclear pore complex protein mAb 414 (obtained from Alexander Palazzo), 1:1000 anti-VDAC1/Porin (Abcam ab110326), 1:1000 anti-Cox2 (obtained from Tom Fox), 1:2500 anti-GAPDH (obtained from Cordula Enenkel), 1:5000 anti-Pgk1 (Abcam ab113687), 1:2000 anti-Killer toxin, and 1:2000 anti-L-A Gag (both obtained from Reed Wickner).

#### Quantification of Nuc1 abundance

Western blots were analyzed with ImageJ analysis software to determine the background-subtracted pixel density of Nuc1-FLAG bands *(6).* To ensure accurate quantification, all blots selected for analysis had similar exposure times. The protein concentration of each fraction loaded for the Western blots was determined using the RD DC Protein Assay (Bio-Rad) and the total protein loaded per lane was derived from this value. Pixel density was normalized to total protein loaded per lane to calculate Nuc1 abundance in each cell fraction.

#### Fluorescent microscopy

Cells were fixed with 4% formaldehyde and washed 3 times with 1x PBS. Fixed cells were then stained with DAPI and mounted on a microscope slide for imaging. Nuc1-GFP images were taken using the Zeiss LSM 880 with Airyscan confocal laser scanning microscope (LSM) on the Fast module and processed with ZEN lite (blue edition) software from Carl Zeiss Microscopy. Por1-GFP/Por2-mCherry images were acquired with the Leica Sp8 confocal LSM and processed with the Leica Application Suite (LAS X) software package. Deconvolution was applied to all images obtained by microscopy.

#### Quantification of colony growth kinetics

Dissections were imaged at 24-hour intervals between 48 to 96 hours post-dissection and the colony sizes of all haploid progeny from tetrad dissections were quantified using ImageJ analysis software (6). The resulting growth curve dataset was plotted to follow the growth kinetics of each spore from the imaged dissections, including nonviable spores with inferable genotypes.

#### Hot acid phenol RNA extraction

Strains were sporulated and harvested by flash-freezing in liquid nitrogen at time points of 0 and 24 hours post-induction. 24-hour samples were subjected to bead-beating (Mini Bead beater, Biospec Products) for 1 minute in Trizol (Invitrogen) followed by 2-minute incubations on ice for 8 cycles. All samples were incubated at 65°C for 30 minutes in phenol solution (Sigma-Aldrich cat# P4682), SDS, and buffer AE (10 mM Tris-Cl, 0.5 mM EDTA pH 9.0) and the phase-separated supernatant was washed with chloroform, precipitated in isopropanol, and washed with 70% ethanol before resuspension in H_2_O.

#### Two-step RT-qPCR

5 micrograms of RNA prepared by the hot acid phenol method was reverse-transcribed using random nonamers and Maxima H Minus Reverse Transcriptase (Thermo Fisher) according to supplier protocols. The resulting cDNA product was isolated by alkaline hydrolysis and RNase A digestion of remaining RNA. Subsequently, qPCR was performed with 1/10 dilutions of cDNA product in 2x SYBR Green/PCR buffer (6) on the iQ5 platform (Bio-Rad). Refer to **Table S4** for qPCR primers used in this study.

#### Normalization and data analysis

The relative starting quantities (SQ) of *M1* and *ACT1* in each sample were determined on the iQ5 software platform (Bio-Rad) using six-point standard curves. *M1* SQs were normalized to endogenous *ACT1* to account for variation in total cDNA concentration between samples. To determine fold change in viral RNA levels, a second normalization to wild-type SQs at 0 hours post-induction was performed for each set of biological replicates.

---

**Figure S1.**
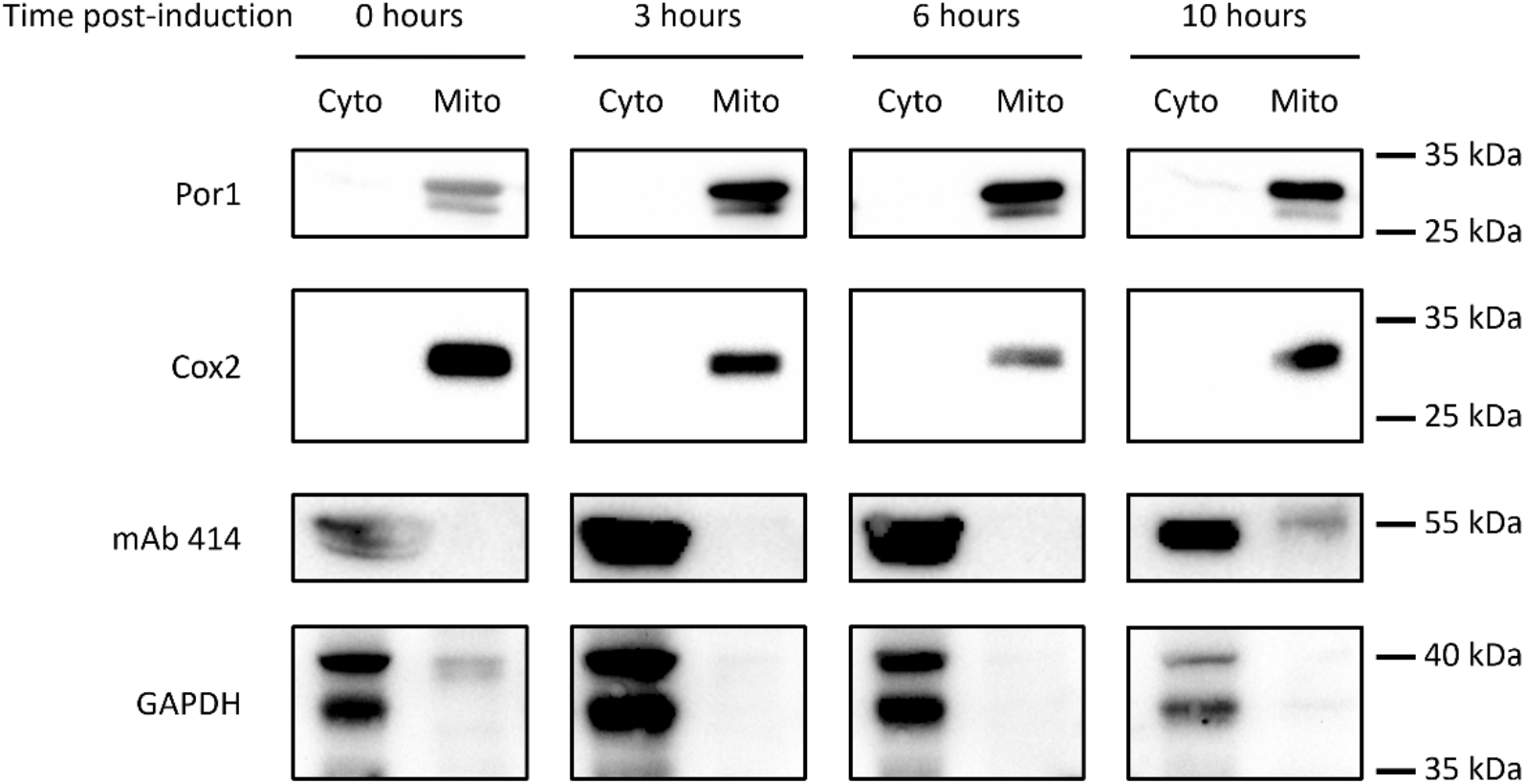
Validation of subcellular fractionation approach. Cell fractions of synchronized meiotic timecourse were probed with antibodies against additional cytosolic (GAPDH) and mitochondrial (Cox2) markers to show appropriate co-fractionation of the selected cell compartment markers. Molecular weight markers are shown on the right.

**Figure S2.**
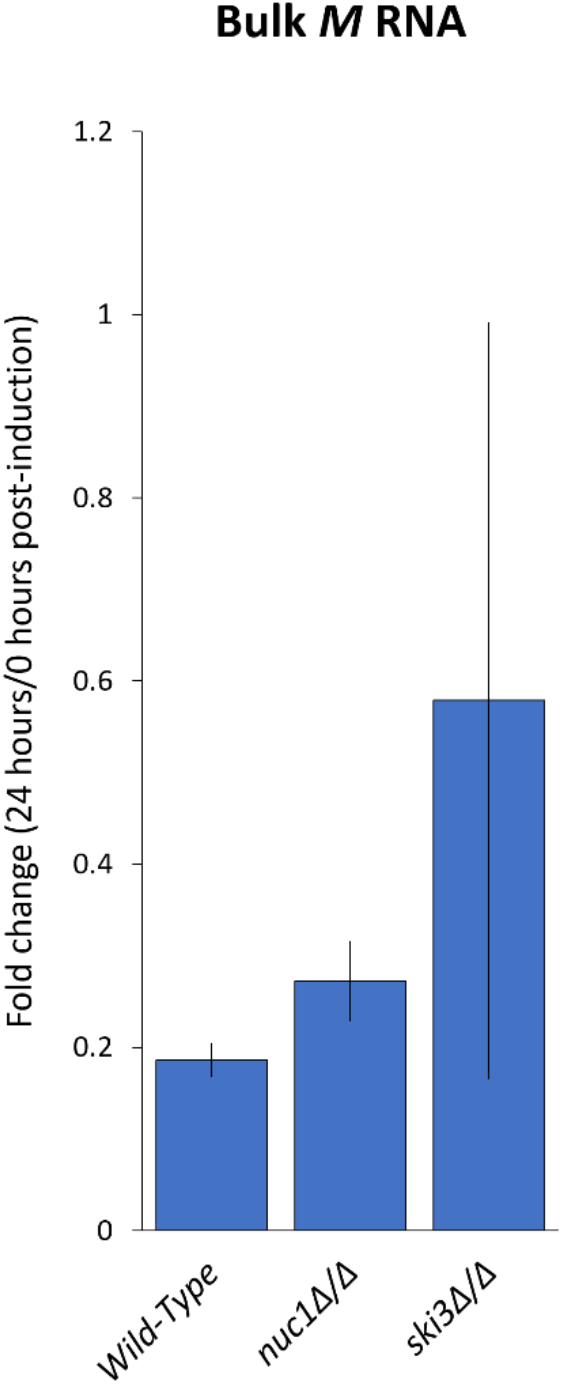
Nuc1 modulates bulk viral dsRNA levels. Bulk *M1* RNA abundance in K^+^ homozygous strains, as quantified by RT-qPCR, in terminally sporulated (24 hours post-induction) populations shown as fold change over pre-meiotic (0 hours post-induction) wild-type *M1* levels. All samples were normalized to endogenous *ACT1* mRNA. Data points represent mean of biological replicates *(n* = 4) ± standard deviation.

**Figure S3.**
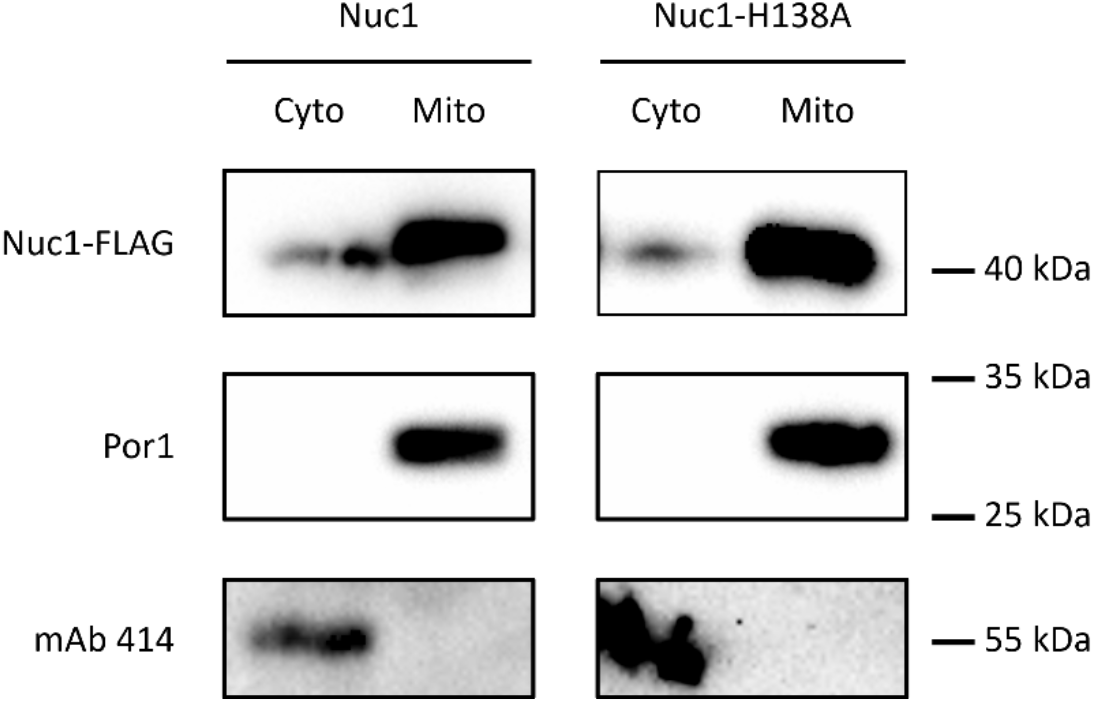
Validation of plasmid-based Nuc1 and Nuc1-H138A constructs. Anti-FLAG Western blot of mitochondrial and cytosolic cell fractions sampled from plasmid-transformed K^+^ *nuc1Δ ski3Δ/+* diploids following induction in SC–HIS+gal media for 6 hours at 30°C to confirm expression of plasmid-based Nuc1 constructs (fractionation controls included). Molecular weight markers are shown on the right.

**Figure S4.**
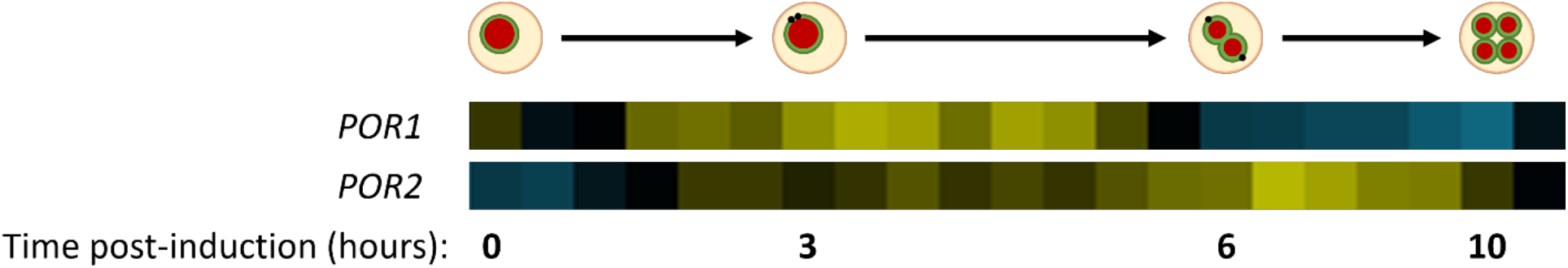
Developmental regulation of *POR1* and *POR2* translation. Meiotic ribosome profiling heatmap of *POR1* and *POR2* derived from the data set provided by Brar et al. (2012) with schematic depiction of meiotic events occurring within the timeframes given. Note that the time elapsed between each data point is not proportional to the physical distances depicted here.

## Supplementary Tables

**Table S1.** Nuc1-mediated viral repression requires nuclease activity. The K^+^ *nuc1Δ/+ ski3Δ/+* parent strain was transformed with plasmids carrying galactose-inducible alleles of FLAG-tagged *NUC1* and dissected on SC-HIS+gal at 30°C. Haploids expressing wild-type Nuc1 or Nuc1-H138A were recovered from multiple independent dissections, genotyped, and counted for the data set. The bottom-most column gives the observed segregation frequency and expected segregation frequency (parenthetical) of the genotype of interest.

|  | **Nuc1** | **Nuc1-H138A** |
| --- | --- | --- |
| K<sup>+</sup> *HIS*<sup>+</sup> *nuc1Δ* *ski3Δ* haploids recovered | 27 | 5 |
| Total K<sup>+</sup> *HIS*<sup>+</sup> haploids recovered | 117 | 120 |
| **Observed frequency of K<sup>+</sup> *HIS*<sup>+</sup> *nuc1Δ* *ski3Δ* haploids** | **0.23 (.25)** | **0.04 (.25)** |

**Table S2.**
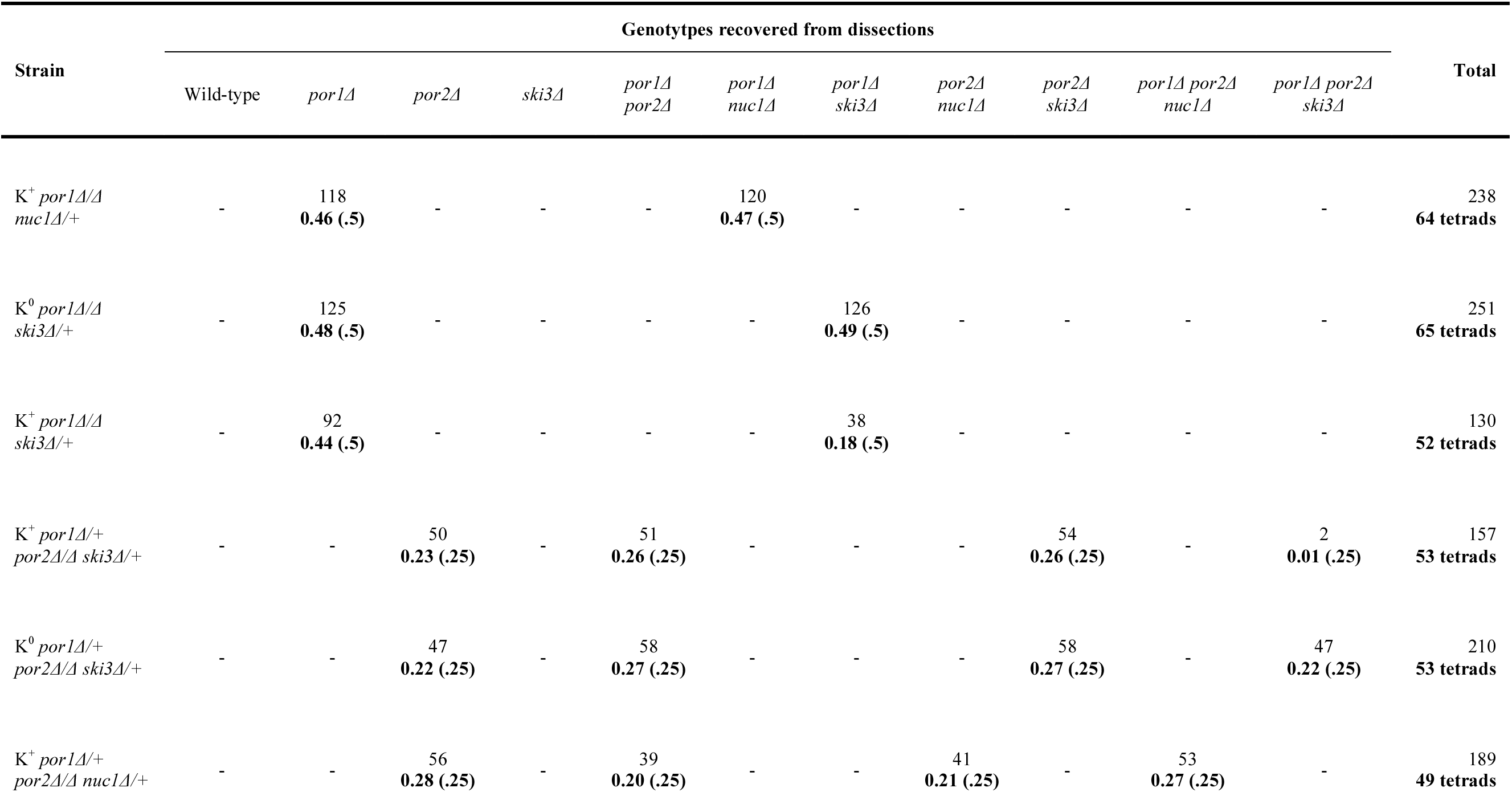
Analysis of *POR1/POR2* genetic interactions. Multiple independent tetrad dissections of porin-*NUC1/SKI3* double and triple mutants were assessed for a binary growth/no growth synthetic lethality phenotype. The total number of haploids of each genotype recovered from each parental genotype is given, with observed segregation frequency (bolded) and expected segregation frequency (bolded, parenthetical) provided below. A dash (-) indicates that no haploids of the given genotype were obtained from dissections of given the parent diploid. The rightmost column gives the total number of spores recovered (top) and number of tetrads analyzed per mutant strain (bottom), which was used to calculate observed segregation frequency.

**Table S3.** Strains list. Numerical identifiers and relevant genotypes of yeast strains used in this study.

| Strain | Relevant Genotype | Plasmids |
| --- | --- | --- |
| HGY432 | W303 *MATa/α K<sup>+</sup> L-A<sup>+</sup> his3/his3*<br>*nuc1::NatMX6/NUC1 ski3::KanMX6/SKI3* | pESC *NUC1-H138A::FLAG HIS3* |
| HGY433 | W303 *MATa/α K<sup>+</sup> L-A<sup>+</sup> his3/his3*<br>*nuc1::NatMX6/NUC1 ski3::KanMX6/SKI3* | pESC *NUC1::FLAG HIS3* |
| MEY322 | SK1 *MATa/α K<sup>0</sup> L-A<sup>+</sup> NUC1-FLAG::KanMX6/NUC1-FLAG::KanMX6* |  |
| MEY783 | SK1 *MATa/α K<sup>0</sup> L-A<sup>+</sup> NUC1-FLAG::LkanMX6/NUC1-FLAG::HygMX6* |  |
| MMY5914 | SK1 *MATa/α K<sup>+</sup> L-A<sup>+</sup>* |  |
| MMY5977 | SK1 *MATa/α K<sup>+</sup> L-A<sup>+</sup> nuc1::NatMX6/NUC1 ski3::HygMX6/SKI3* |  |
| MMY5994 | SK1 *MATa/α K<sup>+</sup> L-A<sup>+</sup> ura3/ura3 nuc1::NatMX6/NUC1 ski3::HygMX6/SKI3* | p5472 *NUC1 CEN/ARS URA3* |
| MMY6106 | SK1 *MATa/α K<sup>+</sup> L-A<sup>+</sup>* |  |
| MMY6117 | SK1 *MATa/α K<sup>+</sup> L-A<sup>+</sup> ski3::HygMX6/ski3::HygMX6* |  |
| MMY6170 | SK1 *MATa/α K<sup>+</sup> L-A<sup>+</sup> nuc1::NatMX6/nuc1::NatMX6* |  |
| MMY6461 | SK1 *MATa/α K<sup>0</sup> L-A<sup>+</sup> nuc1::NatMX6/NUC1 ski3::HygMX6/SKI3* |  |
| MMY6989 | SK1 *MATa/α K<sup>+</sup> L-A<sup>+</sup> por1::KanMX6/POR1 ski3::HygMX6/SKI3* |  |
| MMY7459 | SK1 *MATa/α trp1/trp1 ime1::TRP1/ime1::TRP1 NUC1-FLAG::KanMX6/NUC1-FLAG::KanMX6* |  |
| MMY7464 | SK1 *MATa/α K<sup>0</sup> L-A<sup>+</sup> ski3::HygMX6/ski3::HygMX6* |  |
| MMY7571 | SK1 *MATa/α K<sup>0</sup> L-A<sup>+</sup> por1::KanMX6/por1::KanMX6 NUC1-FLAG::HygMX6/NUC1-FLAG::HygMX6* |  |
| MMY7572 | SK1 *MATa/α K<sup>0</sup> L-A<sup>+</sup> por2::NatMX6/por2::NatMX6 NUC1-FLAG::HygMX6/NUC1-FLAG::HygMX6* |  |
| MMY7599 | SK1 *MATa/α K<sup>+</sup> L-A<sup>+</sup> NUC1-FLAG::KanMX6/NUC1-FLAG::KanMX6* |  |
| MMY7776 | SK1 *MATa/α K<sup>+</sup> L-A<sup>+</sup> nuc1::NatMX6/NUC1 ski3::HygMX6/SKI3 mak3::KanMX6/MAK3* |  |
| MMY7839 | SK1 *MATa/α K<sup>+</sup> L-A<sup>+</sup> nuc1::NatMX6/nuc1::NatMX6 ski3::HygMX6/SKI3 mak3::KanMX6/MAK3* |  |
| MMY7974 | SK1 *MATa/α K<sup>+</sup> L-A<sup>+</sup> nuc1::NatMX6/nuc1::NatMX6* |  |
| MMY7977 | SK1 *MATa/α K<sup>+</sup> L-A<sup>+</sup> ski3::HygMX6/SKI3* |  |
| MMY7978 | SK1 *MATa/α K<sup>+</sup> L-A<sup>+</sup> nuc1::NatMX6/NUC1* |  |
|  | *ski3::HygMX6/SKI3* |  |
| MMY7991 | SK1 MATa/$\alpha$ K<sup>+</sup> L-A<sup>+</sup> *ski3::HygMX6/ski3::HygMX6* | |
| MMY7992 | SK1 MATa/$\alpha$ K<sup>+</sup> L-A<sup>+</sup> | |
| SCY1 | SK1 MATa/$\alpha$ K<sup>+</sup> L-A<sup>+</sup> *por1::KanMX6/POR1*<br>*por2::NatMX6/por2::NatMX6 ski3::HygMX6/SKI3* | |
| SCY164 | SK1 MATa/$\alpha$ K<sup>+</sup> L-A<sup>+</sup> *por1::KanMX6/por1::KanMX6*<br>*ski3::HygMX6/SKI3* | |
| SCY175 | SK1 MATa/$\alpha$ K<sup>0</sup> L-A<sup>+</sup> *por1::KanMX6/por1::KanMX6*<br>*ski3::HygMX6/SKI3* | |
| SCY178 | SK1 MATa/$\alpha$ K<sup>+</sup> L-A<sup>+</sup> *por1::KanMX6/por1::KanMX6*<br>*nuc1::NatMX6/NUC1* | |
| SCY28 | SK1 MATa/$\alpha$ K<sup>0</sup> L-A<sup>+</sup> *por1::KanMX6/POR1*<br>*por2::NatMX6/por2::NatMX6 ski3::HygMX6/SKI3* | |
| SCY41 | SK1 MATa/$\alpha$ K<sup>+</sup> L-A<sup>+</sup> *his3/his3 ura3/ura3*<br>*POR1-EGFP::HIS3/POR1*<br>*POR2-mCherry::URA3/POR2-mCherry::URA3* | |
| SCY53 | SK1 MATa/$\alpha$ *trp1/trp1 ura3/ura3* | pME1 *NUC1::EGFP URA3*<br>pRS424 *ADH-mtRFP TRP1* |
| SCY73 | SK1 MATa/$\alpha$ K<sup>+</sup> L-A<sup>+</sup> *por1::KanMX6/POR1*<br>*por2::NatMX6/por2::NatMX6 nuc1::NatMX6/NUC1* | |

**Table S4.**
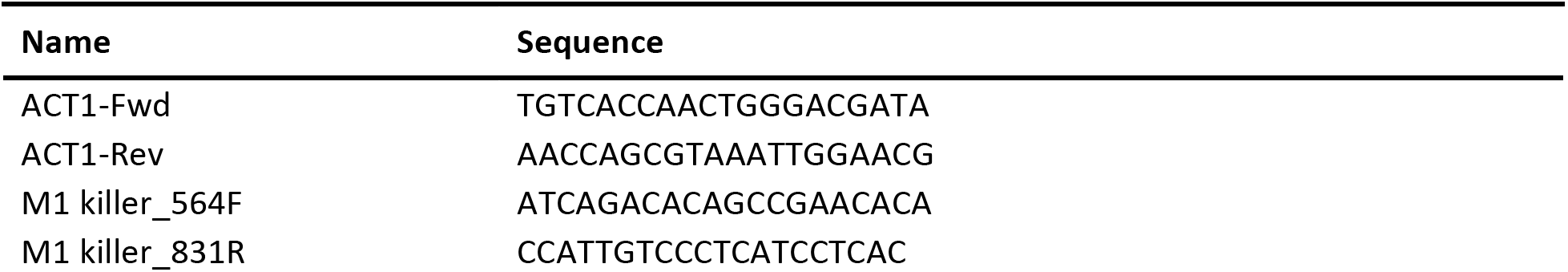
Primer list. Names and sequences of qPCR primers used in this study

